# +microstate: A MATLAB toolbox for brain microstate analysis in sensor and cortical EEG/MEG

**DOI:** 10.1101/2021.07.13.452193

**Authors:** Luke Tait, Jiaxiang Zhang

## Abstract

+microstate is a MATLAB toolbox for brain functional microstate analysis. It builds upon previous EEG microstate literature and toolboxes by including algorithms for microstate analysis for other neuroimaging modalities such as sensor-space MEG and source-space data. +microstate includes codes for performing individual- and group-level brain microstate analysis in resting-state and task-based data including event-related potentials/fields. Functions are included to visualise and perform statistical analysis of microstate sequences, including novel advanced statistical approaches such as microstate-segmented functional connectivity analysis, cluster-permutation topographic ANOVAs, and *χ*^2^ analysis of microstate probabilities in response to stimuli. Additionally, codes for simulating microstate sequences and their associated M/EEG data are included in the toolbox, which can be used to generate artificial data with ground truth microstates and to validate the methodology. +microstate integrates with widely used toolboxes for M/EEG processing including Fieldtrip, SPM, LORETA/sLORETA, EEGLAB, and Brainstorm to aid with accessibility, and includes wrappers for pre-existing toolboxes for brain-state estimation such as Hidden Markov modelling (HMM-MAR) and independent component analysis (FastICA) to aid with direct comparison with these techniques. In this paper, we first introduce +microstate before subsequently performing example analyses using open access datasets to demonstrate and validate the methodology. MATLAB live scripts for each of these analyses are included in +microstate, to act as a tutorial and to aid with reproduction of the results presented in this manuscript.

## 1 Introduction

Magento- and electro-encephalography (M/EEG) are non-invasive tools for functional neuroimaging through recording of extracranial electromagnetic fields generated by electrophysiological cortical activity. M/EEG have been pivotal in uncovering neural mechanisms underpinning healthy cognition and neurological diseases (Lopes da Silva, 2013). In recent years, there has been much interest in the concept of functional brain states, characterised by a discrete (usually small) number of patterns of activation or synchrony across the cortex remaining stable before rapidly transitioning to a different state (Baker et al.,2014; Khanna et al., 2015; Michel and Koenig, 2018; O’Neill et al.,2018).

In the EEG literature, EEG microstate analysis has proven to be a useful tool for studying functional brain states (Michel et al.,2009; Khanna et al., 2015; Michel and Koenig, 2018) in both single-trial data (e.g. resting-state or passive task) (Milz et al., 2016; Seitzman et al., 2017; Michel and Koenig, 2018) and event related potentials (ERPs) (Murray et al., 2008; Koenig et al., 2014). EEG microstate analysis involves clustering spatial topographies of the sensor-space electric potentials (known as ‘maps’) recorded by EEG into a small number of discrete clusters which remarkably typically explain a large amount of variance of the data (Khanna et al., 2015; Michel and Koenig, 2018). The resulting microstate maps are subsequently back-fit to the data, labelling each EEG sample with a microstate label based on maximal similarity to the map in order to obtain a temporal microstate sequence. Microstates have been useful for understanding healthy cognition (Brodbeck et al., 2012; Britz et al., 2014; Milz et al., 2016; Seitzman et al., 2017; Zappasodi et al., 2019), Alzheimer’s disease and other dementias (Nishida et al., 2013; Musaeus et al., 2019; Smailovic et al., 2019; Schumacher et al., 2019; Tait et al., 2020), schizophrenia (Lehmann et al., 2005; Andreou et al., 2014; Tomescu et al., 2014), and other neurological disorders (Khanna et al., 2014)

However, a key limitation of the EEG microstate pipeline is that it is unsuitable for application to MEG or source-reconstructed M/EEG data (Tait and Zhang, 2021). Since MEG has higher spatial resolution than EEG (Boly et al., 2015), and source-reconstruction allows for anatomical interpretation of the electrophysiological data on the cortical level (He et al., 2018), generalization of the microstate pipeline to these modalities is crucial for advancement of understanding the neural mechanisms underpinning brain microstates. Recently, we presented such a generalization of the microstate *k*-means algorithm and applied this algorithm to uncover source-space resting-state microstates and their associations with auditory stimulation (Tait and Zhang, 2021). A number of further advancements were also presented, including validation that microstates were associated with distinct patterns of cortical synchrony and a pipeline to simulate M/EEG sensor- or source-space data with known ground truth microstate maps and microstate sequences (Tait and Zhang, 2021).

Here, we present +microstate, an open-source MATLAB toolbox for multi-modal analysis of microstates in M/EEG sensor- and source-space using the generalized *k*-means algorithm presented by Tait and Zhang (2021). Several open-source toolboxes for sensor-space EEG microstate analysis currently exist such as Cartool, a plugin for the BrainVision Analyzer, and a plugin for EEGLAB (Michel and Koenig, 2018), but +microstate is the only toolbox available for MEG or source-reconstructed microstate analysis. In addition to the generalized *k*-means algorithm for microstate analysis, +microstate can perform a number of other functionalities. While the focus is on *k*-means, +microstate includes options to perform Hidden Markov Modelling (HMM; Baker et al. (2014); Vidaurre et al. (2018)), principal component analysis (PCA), and independent component analysis (ICA; Hyvarinen (1999)) for brain-state estimation using external packages (see subsubsection 2.1.1) in the same framework as our microstate pipeline so that brain-state dynamics using different methodologies can be directly compared and contrasted. Furthermore, +microstate includes functions to calculate microstate-segmented functional connectivity (Hatz et al., 2015, 2016; Tait and Zhang, 2021) which is potentially useful as a tool for studying dynamic functional connectivity beyond the arbitrary sliding window (Tait and Zhang, 2021), as well as perform simulations of data from ground truth microstate sequences (Tait and Zhang, 2021), which is useful for assessing clustering methodologies against a ground truth (Tait and Zhang, 2021). Novel advanced statistical techniques for task-based analysis are additionally included in the toolbox, including cluster-permutation (Maris and Oostenveld, 2007) extensions to topographic ANOVAs (TANOVAs) (Murray et al., 2008) and *χ*^2^ statistics of microstate probabilities in response to stimuli (Tait and Zhang, 2021). +microstate is based around objects which store the data and have intuitively named functions for analysis, simulation, visualization, and statistics, resulting in a simple, accessible toolbox for microstate analysis with limited coding background required.

This manuscript is structured as follows. In Materials and Methods, we outline the requirements, implementation and usage of +microstate, including key commands and how data is stored. In Results, we demonstrate the results of some examples of usage with empirical/simulated, sensor/source-space M/EEG data, particularly with a focus on benchmarking against ground truths to validate results are as expected. This includes analysis of resting-state EEG to reproduce the well established canonical EEG microstates (Michel and Koenig, 2018), analysis of MEG event related fields (ERFs) to uncover a sensor-MEG microstate associated with different levels of activation under different experimental conditions, analysis of source-space MEG microstate sequences in response to auditory stimuli, and reproducing ground truth source-space microstate maps and microstate statistics in a simulated experiment. Codes and data for these analyses are freely available and included in +microstate as MATLAB Live Script tutorials, and the figures reported here were produced using MATLAB’s default random number generator seed to increase transparency and reproducibility. Therefore the examples presented here may act as validation of the toolbox, examples of possible use cases of the toolboxes, and as a tutorial.

## 2 Materials and Methods

### 2.1 Installing the toolbox

+microstate can be freely downloaded from https://github.com/lukewtait/microstate_toolbox. To use the toolbox, MATLAB should be installed and the folder +microstate should placed into a folder on the MATLAB path.

#### 2.1.1 Requirements

+microstate is compatible with MATLAB R2017b and later versions; rigorous testing with earlier versions of MATLAB have not been performed. A number of MATLAB built in toolboxes are required for full functionality, including the Statistics and Machine Learning toolbox, Signal processing toolbox, and Wavelet toolbox (only required for simulations of random walk sequences). A number of external toolboxes are also used and the required functions are included with +microstate under `microstate.external`. These include FastICA v2.5 (https://research.ics.aalto.fi/ica/fastica/) to use ICA for clustering, HMM-MAR (https://github.com/OHBA-analysis/HMM-MAR) to use Hidden Markov Modelling for clustering, and freely available custom-written scripts for data visualization of summary statistics (https://github.com/lukewtait/matlab_data_visualization).

### 2.2 The ‘individual’ object

All data from a given participant or scan is stored in an object of the *individual* class. Methods for this class include the functionality to import and preprocess the data, perform individual-level microstate analysis, calculate statistics of microstate sequences, and calculate microstate-segmented functional connectivity. An empty *individual* object can be created by calling `ms = microstate.individual` in the command line, where `ms` is a microstate individual object. The electrophysiological data is stored in the class properties:

- `data`: A *T* × *N* double array storing the MEG/EEG/source reconstructed time series, where *T* is the number of samples and *N* is the number of sensors/ROIs.
- `time`: A vector of length *T* containing the time axis for the data.
- `modality`: A character array, which can be either `‘meg’, ‘eeg’, ‘source’`, or `‘ampenv’` depending on the modality of the data.
- `bad_samples`: A vector containing the indices of bad samples, such as samples which contain artifacts.

Once microstate analysis has been performed, other relevant properties of the class will be filled, such as:

- `gfp`: A vector of length *T* containing the GFP of each sample.
- `maps`: A *N* × *k* double array storing the microstate maps resulting from cluster analysis, where *k* is the number of clusters.
- `label`: A vector of length *T* containing the integer microstate label of each sample resulting from cluster analysis.
- `stats`: A structure which can output statistics of the microstate sequences.
- `networks`: A cell array of length *k* containing *N* × *N* functional networks.

A full list of properties and methods for the *individual* object can be found by typing help `microstate.individual` in the MATLAB command line.

#### 2.2.1 Adding and preprocessing data

To construct a non-empty microstate *individual* object, the following command is used:

~~~
ms = microstate.individual(data,modality,time) ;
~~~

where the inputs `data, time`, and `modality` fit the descriptions of the class properties with the same name given above (note, alternatively `time` can be given as a single value corresponding to a sampling rate, and the time axis will be automatically generated). Bad samples, for example those identified by artifact detection algorithms, can be added by typing `ms = ms.add_bad_samples(bad_samples)`.

Alternatively, an empty *individual* object can be created as `ms = microstate.individual`, and data can be subsequently be added. If data is in the correct format for +microstate, it can be added using the `add_data` function. Additionally, +microstate integrates with a number of other M/EEG processing toolboxes, allowing for data to be imported directly from these toolboxes. The toolboxes integrated with +microstate (and their respective +microstate functions to import data) are:

- Fieldtrip (Oostenveld et al. (2011); http://fieldtriptoolbox.org). Import data using the function `ms = ms.import_fieldtrip(data)`, where `data` is a Fieldtrip raw, timelock, or or source structure.
- SPM (https://www.fil.ion.ucl.ac.uk/spm/). For MEEG objects, use the function `ms = ms.import_spm_meeg(D)`, where `D` is the SPM MEEG object. For source data, one must first calculate the inverse filter in SPM (automatically saving the filter in the MEEG object), and then call the function as `ms = ms.import_spm_meeg(D, ‘source’)`. Additionally, 4D volumetric SPM images saved as NiFTi files can be imported by calling `ms = ms.import_spm_nifti(filename,fsample)`, where `filename` is the path to the NiFTi file and fsampie is the sampling rate of the data.
- EEGLAB (Delorme and Makeig (2004); https://eeglab.org). `ms = ms.import_eeglab(EEG)`, where `EEG` is an EEGLAB data object. Note that EEGLAB integrates with Fieldtrip for source reconstruction, so only sensor-space data can be called using `import_eeglab`.
- LORETA and sLORETA/eLORETA (http://www.uzh.ch/keyinst/loreta). Import source reconstructed data using the command `ms = ms.import_loreta(filename,fsample)`, where `filename` is the path to the LORETA binary (*.lorb) or sLORETA binary/text (*.slor/*.txt) file and `fsample` is the sampling rate of the data.
- Brainstorm (Tadel et al. (2011); http://neuroimage.usc.edu/brainstorm). For sensor data, call `ms = ms.import_brainstorm(bst_data,modality)`. For voxel-wise source data, use `ms = ms.import_brainstorm(bst_data,bst_source)`. For parcellated/ROI source data, call `ms = ms.import_brainstorm(bst_data,bst_source,bst_scouts)`. Descriptions of the structures bst_data, bst_source, and bst_scouts and how to export them from Brainstorm to MATLAB can be found in the import_brainstorm function help.

While the focus of +microstate is on microstate analysis as opposed to data preprocessing, some simple preprocessing commands are included. Functions for preprocessing include re-referencing EEG data to average, resampling, bandpass filtering, orthogonalization (Colclough et al., 2015), and calculating the amplitude envelope. A typical microstate pipeline might include bandpass filtering 1-30 Hz, re-referencing to average (EEG only), and resampling to 256 Hz (if data is sampled at faster rates) (Tait and Zhang, 2021). This default pipeline is simple to call and implemented as a default, by calling

~~~
ms = ms.preprocess ;
~~~

Other functions are called in a similar manner, e.g. to bandpass filter in the range `a-b` Hz, one could call `ms = ms.preprocess_filter(a,b)`. Almost all functions have optional inputs which can be called in name-value pairs, e.g. for a band-stop filter between 48-52 Hz to attenuate line noise, type

~~~
ms = ms.preprocess_filter (48,52, ‘type’,’stop’) ;
~~~

#### 2.2.2 Performing microstate analysis using k-means clustering

A full description of the generalized modified *k*-means algorithm is given in Tait and Zhang (2021). Briefly described, the algorithm involves the following steps. Firstly, the data must be appropriately transformed depending on whether it is EEG (re-referenced to average), MEG (no transform), source (absolute value taken), or amplitude envelope data (no transform). These transforms are automatically handled within the functions of the toolbox based on the `modality` property of the individual object, and do not need to be performed manually by the user at any stage. The GFP for a given sample *t* is given by

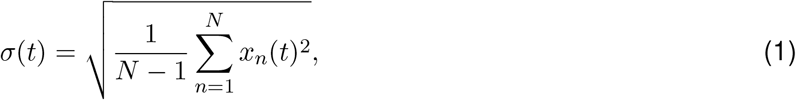

where *x_n_*(*t*) is the value of the *n*’th sensor/ROI at the *t*’th sample. For EEG data, this is the standard deviation (due to average reference). For all other datasets, it can be viewed as a normalized vector norm. In both cases, peaks of the GFP correspond to samples with local maxima of signal-to-noise ratio and topographic stability (Koenig and Brandeis, 2016; Tait and Zhang, 2021). In +microstate, the GFP can be calculated by calling `ms = ms.calculate_gfp`, and plotted by calling `ms.plot(‘gfp’)`.

Subsequently, GFP peaks are extracted and a generalized version of the modified *k*-means algorithm presented by Pascual-Marqui et al. (1995) is run. This algorithm is equivalent to a typical *k*-means clustering algorithm, but instead of Euclidean distance, map dissimilarity is used. Map *similarity* between two spatial topographies **x** ∈ *N* × 1 and **y** ∈ *N* × 1 is given by

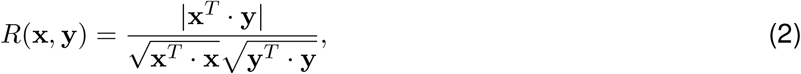

and the map *dissimilarity* (or distance) is given by 1 –*R*(**x, y**). The command below runs the *k*-means clustering in +microstate,

~~~
ms = ms.cluster_estimatemaps(k)
~~~

where `k` is the number of clusters. The toolbox uses kmeans++ to choose initial maps for clustering (Arthur and Vassilvitskii, 2007), and by default uses a maximum of 100 iterations and 20 replicates, although these values can be changed using the name-value pair inputs `‘kmeans_maxiter’` and `‘kmeans_replicates’` respectively. After running this function, the *individual* object ms will have non-empty properties maps and `label`, corresponding to the microstate maps and temporal microstate sequence respectively.

The choice of *k*, the number of states, is a free parameter. To use an algorithm to identify the optimum number of states, the function `cluster_koptimum` can be called as follows

~~~
ms = ms.cluster_koptimum ;
~~~

By default, this will run the clustering analysis for 2-20 states and use the kneedle algorithm (Satopää et al., 2011; Tait and Zhang, 2021) to select an optimum value. Other choices of criterion can be selected using the name-value pair input criterion, which can take values of `‘KrzanowskiLai’` (Murray et al., 2008), `‘CrossValidationIndex’` (Pascual-Marqui et al., 1995), or the four available criteria in the MATLAB function `evalclusters` (https://www.mathworks.com/help/stats/evalclusters.html#shared-criterion). Other name-value pair inputs include `‘kmin’` and `‘kmax’` which take integer values and control the maximum and minimum values of *k* to search respectively. The object `ms` will be updated to contain the microstate maps and temporal labels for the optimum number of microstates. By calling additional outputs to this function, the microstate maps and labels for all values of *k* can be saved.

#### 2.2.3 Performing microstate analysis using other algorithms

+microstate also includes options to perform microstate analysis using PCA, ICA, and HMM by integrating with external toolboxes. It should be noted that for these algorithms, the functions `cluster_estimatemaps` and `cluster_koptimum` simply act as wrappers for these external toolboxes to give outputs in the same format as the *k*-means output. These alternative methods can be called in the clustering functions described above using the name-value pair input `‘clustermethod’`, with value `‘pca’`, `‘ica’`, or `‘hmm’` respectively. PCA uses the `pca` function in the MATLAB Statistics and Machine Learning Toolbox. ICA uses the fastica algorithm (https://research.ics.aalto.fi/ica/fastica/). HMM uses the HMM-MAR toolbox (https://github.com/OHBA-analysis/HMM-MAR), by default using the standard options from the example scripts in this toolbox which are based on the pipelines of Baker et al. (2014) for amplitude envelope data and Vidaurre et al. (2018) for all other modalities. Other options can be included by specifying the name-value pair input `‘hmm’` which contains the options structure used when calling the HMM-MAR toolbox (see the documentation for this toolbox for more details). For PCA and ICA, any of the criterion of choosing number of states described above can be used. For HMM, the value of *k* which minimises free energy is used.

#### 2.2.4 Analysing microstate sequences

Once clustering analysis has been performed as described above, a range of statistics of the microstate sequences can be calculated. Global statistics of the microstate sequences include GEV (Murray et al., 2008), mean duration of microstates (Koenig et al., 2002), Hurst exponent of the sequences (Van De Ville et al., 2010), microstate complexity (Tait et al., 2020; Tait and Zhang, 2021), and the autoinformation function of the microstate sequence (von Wegner et al., 2017). Class-specific statistics include mean duration of the microstates within a particular class (Koenig et al., 2002; Lehmann et al., 2005), coverage of a class (the percentage of time spent within a class) (Lehmann et al., 2005), and occurrences of the class (number of times the state appears per second) (Lehmann et al., 2005). +microstate can also calculate the Markov and syntax matrices (with and without self-transitions respectively) (Lehmann et al., 2005; Nishida et al., 2013; von Wegner et al., 2017), the information-theoretical zeroth and first order Markov statistics and their *p*-values (von Wegner et al., 2017), and test for non-random microstate syntax (Lehmann et al., 2005; Nishida et al., 2013).

Each statistic is given by a function beginning with `stats_`, e.g. to calculate GEV you can call the function `stats_gev.` +microstate also includes a wrapper function for each of these stats functions, which can be called as

~~~
ms = ms.stats_all ;
~~~

The property `stats` of the object `ms` will be updated to include values for each of the stats described above.

#### 2.2.5 Microstate-segmented functional connectivity

Microstate-segmented functional connectivity patterns (Hatz et al., 2015, 2016; Tait and Zhang, 2021) can also be calculated in +microstate. This can be performed by calling

~~~
ms = ms.networks_wpli(freq_band) ;
~~~

While the input `freq_band = [f_low,f_high]` is optional, phase synchrony is typically calculated from narrowband signals and hence it is strongly recommended to include this input in order to specify a frequency band (from `f_low` to `f_high` Hz, e.g for the 8-13 Hz alpha band specify `freq_band = [8,13]`) in which to calculate connectivity.

#### 2.2.6 Visualizing microstate data

+microstate includes options for visualizing the data and microstate sequences stored in the *individual* object using the `plot` function. This can be called as follows

~~~
ms.plot (string) ;
~~~

Here, `string` is a string specifying what should be plotted, and can take on a wide range of values. For example, to plot the electrophysiological timeseries, you can call `ms.plot(‘data’)`. There are options to plot many other statistics including GFP, power spectrum, microstate maps, Markov/syntax matrices, autoinformation functions, coverage/duration/occurrence of microstate classes, and microstate segmented functional connectivity patterns.

### 2.3 The ‘cohort’ object

While the *individual* object is useful for performing microstate analysis at the level of an individual participant or scan, the *cohort* object can be used for group level analysis. An empty *cohort* object can be created by calling `coh = microstate.cohort` in the command line, where coh is a microstate cohort object. Properties of this class include:

- `individual`: An array of *M individual* objects, where *M* is the number of participants/scans and each element in the array corresponds to a participant/scan.
- `condition` and `conditionlabels`: These values are used when the data contains multiple conditions (e.g. multiple scans, disease vs control groups, experiment vs rest, etc) to specify which *individual* object belongs to which condition. `condition` is an array of length *M* taking on integer values from one to the number of conditions, while `conditionlabels` gives labels for each condition. For example, if `coh.conditionlabels = {‘task1’,’task2’,’ rest’}` and `coh.condition(1)=2`, then `coh.individual(1)` was recorded during the *task2* condition.
- `globalmaps`: A *N* × *k* double array storing the group level microstate maps resulting from global clustering.
- `stats`: A structure which can output statistics of the microstate sequences
- `process`: A table containing a record of all processes performed on the data.

#### 2.3.1 Adding individuals to a cohort

Let `ms` be a microstate *individual* object and coh be a *cohort* object. The `add_individuals` property of the *cohort* class can be used to add the individual as follows:

~~~
coh = coh.add_individuals(ms) ;
~~~

Now, let us assume we have two *individual* objects recorded during two conditions *condition1* and *condition2*. We can add these to a cohort object as

~~~
coh = coh.add_individuals(ms1,’condition1’) ;
coh = coh.add_individuals(ms2,’condition2’) ;
~~~

For large cohorts, including all of the data can be very memory intensive. It therefore might be preferable to take more memory efficient approaches if not all the data is required. For example, one could read in each dataset individually, perform clustering on that dataset, and then to obtain group level maps perform clustering on the sets of individual maps (Khanna et al., 2014), in which case no data (only the individual maps) would be required for clustering. Alternatively, one might sample a subset of GFP peaks from each individual for clustering (Tait and Zhang, 2021). A third input to the `add_individuals` function allows for specification of how much data is stored; a value in the range 0-1 specifies to store a random fraction of the data (e.g. 0 stores none of the data, 0.1 would randomly select 10% of data points to store, and 1 stores all data), while integer values greater than 1 specify a fixed number of GFP peaks to store.

#### 2.3.2 Calling ‘individual’ functions for all individuals within a cohort

Many of the functions that apply to the *individual* class can be called for the *cohort* class with the prefix `ind_`. Any function with this prefix is simply a wrapper, looping over all individuals in the cohort. For example, calling `coh = coh.ind_preprocess_filter(1,30)` is equivalent to running a for loop through all *individuals* in the *cohort* and using the `preprocess_filter` function to bandpass filter 1-30 Hz.

#### 2.3.3 Performing group-level microstate analysis

Group level clustering can be performed using the functions `cluster_global` and `cluster_globalkoptimum`, which are group level equivalents to the *individual* class functions `cluster_estimatemaps` and `cluster_koptimum` respectively. In fact, these functions generate a new *individual* object whose data property is a concatenation of all stored data points from all individuals and calls the individual clustering functions on this concatenated data set. To perform group level clustering, call

~~~
coh = coh.cluster_global (k) ; % option 1: cluster k maps
coh = coh.cluster_globalkoptimum ; %option 2: find optimum k and cluster
~~~

These commands will update the property globalmaps of the *cohort* object to include the clustered maps.

To first perform clustering analysis on the individual level and then cluster the individual maps (Khanna et al., 2014), you can call the following:

~~~
coh = coh.ind_cluster_estimatemaps(k) ;
coh =coh.cluster_global(k,’cohortstat’,’maps’) ;
~~~

#### 2.3.4 Performing trial-based ERP/ERF microstate analysis

The *cohort* object can also be used to store multiple trials under multiple conditions for one or more participants. This is particularly useful for analysis of ERPs/ERFs (Murray et al., 2008; Koenig et al., 2014). In this case, you can perform permutation tests on GFPs or topographies for both between-trial or within-participant designs, using the methodologies outlined by Murray et al. (2008). For example, to perform TANOVA tests, we can call

~~~
[p, stats] = coh.erp_clusterperm_TANOVA;
~~~

The millisecond-by-millisecond TANOVA results (Murray et al., 2008) are stored in the output `stats.p_sample`. However, this approach has the issue of needing to correct for multiple comparisons (i.e. one p-value per sample of data), and hence +microstate also implements a cluster permutation TANOVA combining the map dissimilarity (DISS) statistic (Murray et al., 2008) and permutation methodology from millisecond-by-millisecond TANOVAs with the multiple comparison cluster approach used in cluster permutation testing (Maris and Oostenveld, 2007). That is, DISS is calculated and condition labels are permuted in the same manner as described for the millisecond-by-millisecond TANOVAs (Murray et al., 2008), but instead of comparing empirical DISS against the distribution of permuted DISS to obtain a *p*-value at each time point, we choose a threshold (here the midway point between maximum and minimum DISS values), find clusters of neighbouring points exceeding the threshold, and calculate the sum of DISS over all samples within a cluster to obtain a cluster DISS value. We subsequently compare the maximal empirical cluster statistic against the null maximal cluster statistics to obtain a multiple-hypothesis corrected *p*-value for the cluster, which is reported in the output `p`.

Similarly, one can calculate the *χ*^2^ distance between the histograms of microstate probabilities in two sets of conditions and perform cluster permutation analysis by calling

~~~
stats = coh.erp_chi2stateprobs(expectedConditions,observedConditions) ;
~~~

where `expectedConditions` and `observedConditions` are the names of the conditions to be compared. For example, in subsection 3.4 we compare standard and deviant stimuli, so treat the standard stimuli as the expected (or baseline) probabilities of microstate classes, and the deviants as the observed probabilities. Alternatively, one can specify the expected conditions to be `‘prestim’` to compare pre- and post-stimulus periods as presented in Tait and Zhang (2021), but in this case cluster permutation analysis is not available and only millisecond-by-millisecond *p*-values are returned.

### 2.4 Simulations

Tait and Zhang (2021) presented a methodology for simulating microstates. This takes on three steps, which are easily implemented in +microstate. Firstly, a microstate sequence must be simulated. Two options are implemented in +microstate, including a random walk decision tree (Tait and Zhang, 2021) or simulating from a pre-specified Markovian transition matrix. These simulated sequences are generated by making an *individual* object and then calling functions `simulate_seq_randomwalk` or `simulate_seq_markov`, which will update the `label` property of the *individual* object. Secondly, microstate maps must be added. This is as simple as specifying the `maps` property of the *individual* object. Thirdly, the data must be simulated. This is done by calling the `simulate_data` function, which will update the *individual* object’s `data` property to include the simulated MEG/EEG/source data. By default, the parameters of the neural mass model are those previously reported (Deco et al., 2009; Abeysuriya et al., 2018; Tait and Zhang, 2021), but the parameters can be customized using the name-value pair input `‘params’`, which takes a value of a structure specifying any custom parameters.

Alternatively, simulations can be called via the wrapper function

~~~
ms = ms.simuiate ;
~~~

which performs all steps of this pipeline.

## 3 Results

In this section, we will demonstrate some examples of using +microstate for microstate analysis of real data and simulations. In the first and second examples, we will use +microstate to perform sensor-level microstate analysis on free open access EEG and MEG datasets. Subsequently, we will simulate an example experiment and demonstrate group level analysis. The aims of these examples are to test against known benchmarks to validate the toolbox. In the first example (resting-state EEG data), we aim to recreate the four canonical resting-state EEG microstate maps (Michel and Koenig, 2018). In the second example (MEG evoked fields), we use a dataset where known topographical differences between ERFs are exhibited at a particular latency. We test a hypothesis that these topographical difference are related to activation of a particular microstate class with a similar topography during this latency. In the third example (simulated source-space data), we aim to test whether the microstate pipeline can recreate ground-truth group-level differences between simulated cohorts. In the fourth example, we analyse microstate sequences from source-space MEG recordings during an auditory task, as presented in Tait and Zhang (2021). For each of these examples, all data and a MATLAB Live Script tutorial is included with the toolbox such that all figures and results can be reproduced exactly. Finally, we will briefly give an overview of the results of Tait and Zhang (2021) when applying the toolbox to source-reconstructed resting-state MEG data.

### 3.1 Recreating the canonical EEG microstate maps

EEG microstate analysis has widely been applied to resting-state data, identifying an optimum of four states known as the canonical microstates (Michel and Koenig, 2018). When the data is sensor-space EEG, the generalized formalism of the microstate algorithm implemented in +microstate is identical to the widely used EEG microstate modified *k*-means algorithm. Hence, +microstate can be used for traditional EEG microstate analysis as well as MEG or source space microstate analysis. In this section, we demonstrate an example of how to perform EEG microstate analysis using +microstate.

The dataset used here is the open-access data supplied with the Fieldtrip tutorial for cleaning and pre-processing resting-state data (https://www.fieldtriptoolbox.org/workshop/madrid2019/tutorial_cleaning/), and is described in Chennu et al. (2016). We downloaded the data set and pre-processed the data according to this webpage. For efficiency, we also downsampled the data to the 20 electrodes which correspond to the 10-20 system, since this is sufficient for accurate estimation of microstates (Khanna et al., 2014). +microstate was used to preprocess the data (re-reference to average and 1-30 Hz bandpass filter) and perform cluster analysis for 4 microstates. Figure 1 shows the resulting microstate maps, plotted using +microstate’s `plot(‘maps’)` function. The resulting maps closely correspond to the four canonical microstate maps (Michel and Koenig, 2018). Analysis of this microstate sequence in +microstate demonstrates microstate statistics are within ranges reported in the literature, including a mean duration of 54 ms (von Wegner et al., 2017; Tait et al., 2020), and GEV of 55% (Michel and Koenig, 2018), and non-Markovian transitioning indicated by a highly significant information-theoretical Markov statistic (von Wegner et al., 2017) and a Hurst exponent of 0.60 (Van De Ville et al., 2010; von Wegner et al., 2016, 2018).

**Figure 1:**
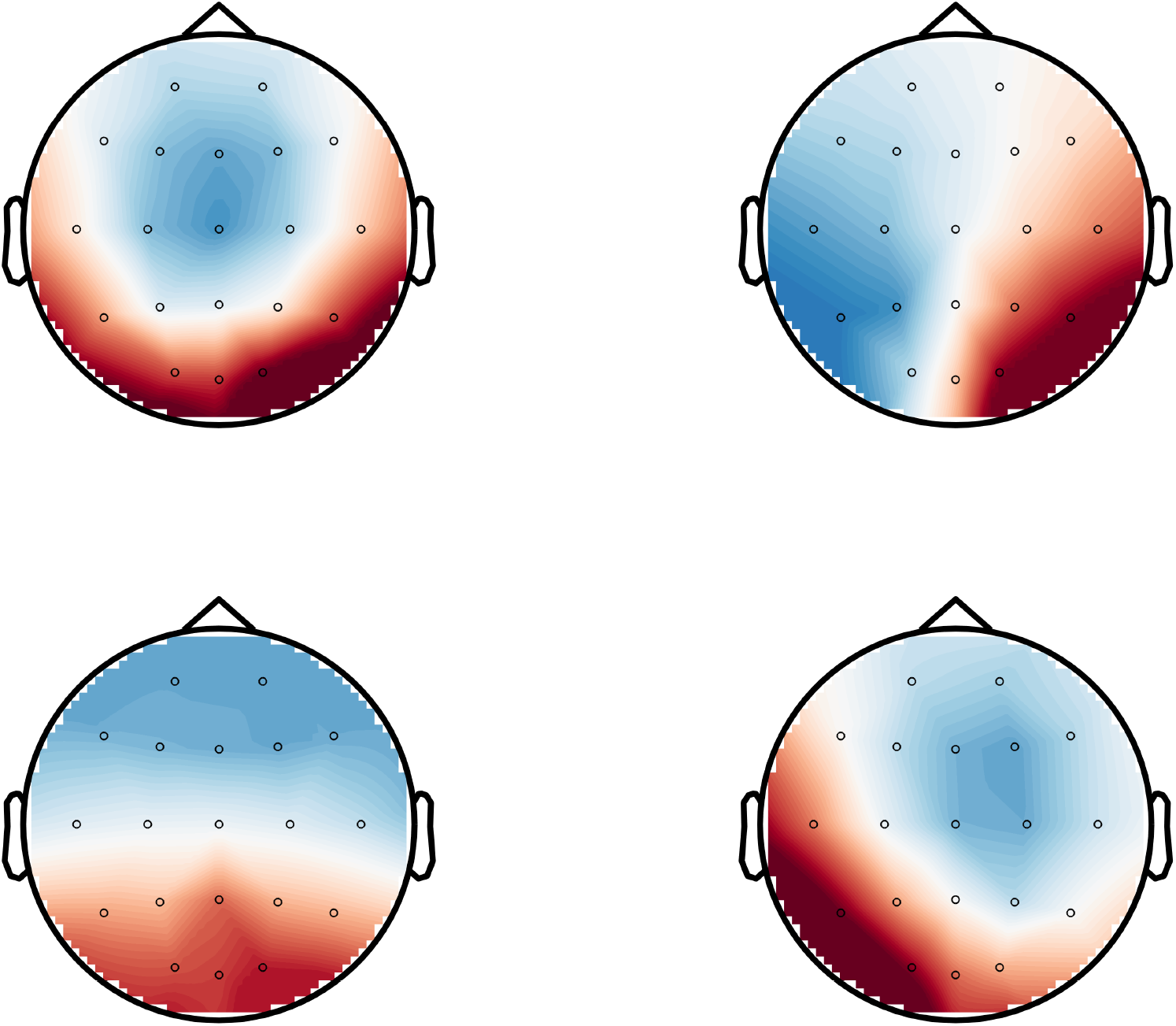
EEG microstate maps. derived from a single-subject open-access resting-state EEG scan using the +microstate toolbox and plotted using the toolbox’s `plot (‘maps’)` function. The maps closely correspond to the canonical maps A (top right), B (bottom right), C (bottom left), and D (top left) (Michel and Koenig, 2018).

### 3.2 Topographic microstate analysis of MEG ERPs

Murray et al. (2008) proposed a topographic and microstate methodology for analysing event-related potentials (ERPs). In this section, we used +microstate to perform topographic microstate analysis on the open-access MEG ERP data supplied with the Fieldtrip tutorial for cluster-based permutation tests on event-related fields (https://www.fieldtriptoolbox.org/tutorial/cluster_permutation_timelock/). This dataset is from a study in which MEG was recorded following congruent and incongruent sentences (Wang et al., 2012). The data was downloaded and pre-processed according to the Fieldtrip tutorial. Following the Fieldtrip tutorial, we analysed data in the range 0-1 seconds following the stimulus.

Here, we will demonstrate results from between-trial experimental design comparing congruent and incongruent trials between participants, but +microstate can also be used for within-subject designs with only minor adaptions to the script (an example script is included in the toolbox). In the Fieldtrip tutorial, cluster permutation testing on the ERP time courses demonstrated significant differences between congruent and incongruent trials 550-750 ms following, localized to left frontal and parietal electrodes. The topography of these differences are shown in Figure 2A. In this section, we hypothesised there is a cortical network (or microstate) associated with this topography which differs in activation between congruent and incongruent trials, and used +microstate to test this hypothesis.

**Figure 2:**
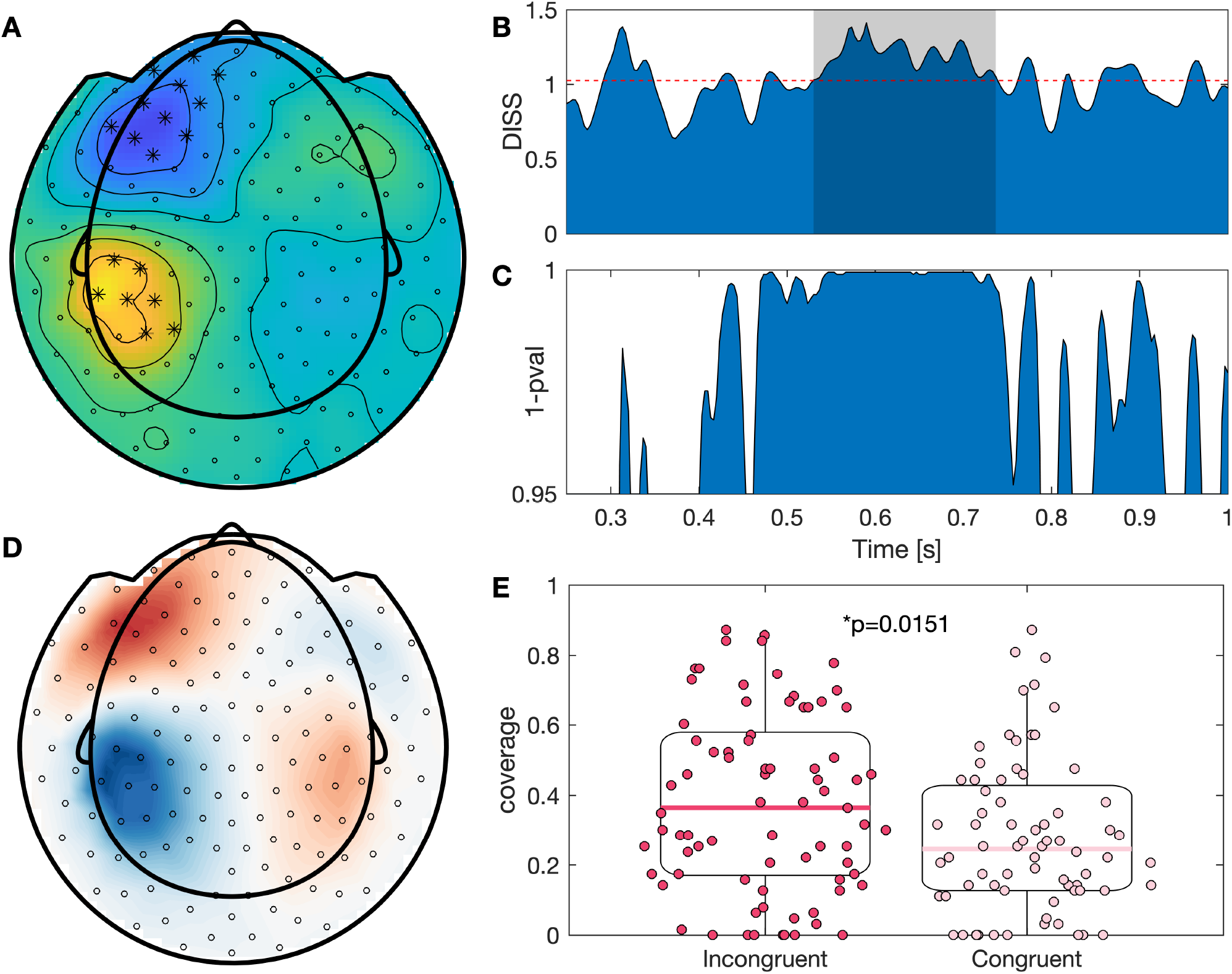
Topographic microstate analysis of MEG ERPs. (A) Result of analysing the data in Fieldtrip following the cluster permutation analysis tutorial (see text for description) between 550-750 ms, and plotted using Fieldtrip. Shown is the difference between congruent and incongruent responses. Sensors marked with an asterisk were significantly different between conditions. (B) Map dissimilarity (DISS) between congruent and incongruent trials. The horizontal dashed line shows the threshold used for cluster permutation TANOVAs, and the shaded region shows the significant clusters from cluster permutation TANOVAs. The timing of significant differences are between 530-737 ms. (C) In contrast to cluster permutation TANOVAs, which correct for multiple comparisons, here we plot millisecond-by-millisecond TANOVA results without correction for multiple comparisons following Murray et al. (2008). (D) One of the four microstate maps identified from clustering analysis, and plotted using +microstate. Due to the correspondence between this microstate and the difference map shown in A, we hypothesise this microstate is associated with a network which differs in activation between congruent and incongruent trials. (E) There is a significant difference between congruent and incongruent trials in the coverage of the microstate shown in D between 530-737 ms.

Firstly, topographic ERP analysis (Murray et al., 2008) was used to find time samples with significant differences between the conditions. Figure 2B shows the global map dissimilarity between the grand average ERPs for each condition. Murray et al. (2008) proposed a millisecond-by-millisecond TANOVA approach to identify samples in which the topography differs between conditions. Results of millisecond-by-millisecond TANOVA are shown in Figure 2C. However, this approach has the issue of needing to correct for multiple comparisons, so here we instead analyse the output of the cluster permutation TANOVA approach (subsubsection 2.3.4). Figure 2B shows that a significant cluster was identified between 530-737 ms using this cluster permutation TANOVA approach, in line with the differences observed for classical ERP analysis in the Fieldtrip tutorial.

We subsequently performed microstate analysis on the ERPs as described in Murray et al. (2008) and implemented in +microstate. We used the Krzanowski-Lai criterion for choosing the number of states, restricting this to greater than three states (Murray et al., 2008). Four states were optimum. Of these four states, one showed close correspondence with the ERP difference map (Figure 2D), supporting our hypothesis that there is a cortical network which is associated with differences in response between congruent and incongruent trials. To further test the hypothesis that this network differs in activation between conditions, we quantified the coverage of this state between 530-737 ms, and found a significantly different coverage (*p* = 0.0151, Wilcoxon-rank sum test; Figure 2E).

### 3.3 Simulations and group-level analysis of cohorts under different conditions

In this section, we demonstrate simulations and group-level analysis of cohorts in +microstate. The simulated experiment is as follows: We have *n* = 20 participants, who undergo two M/EEG scans. In the first scan, conditions are such that the microstate sequence is generated by a random walk decision tree, and hence should exhibit long range temporal correlations. In the second scan, the microstate sequence is generated by a Markov chain; specifically a Markovian surrogate to the random walk sequence (i.e. coverages and syntax are theoretically identical to the random walk sequence, but no long range correlations exist). All other parameters are equal between conditions, e.g. microstate maps and equations for neural dynamics. In the first section below, we will describe how to simulate such an experiment, generating artificial data and creating a +microstate *cohort* structure. In the subsequent section, we will treat the simulated data as if it were real data from an experiment and perform microstate analysis, making the hypothesis that the estimated microstate sequences should demonstrate evidence of more Markovian properties in the second condition.

#### 3.3.1 Simulating the experiment

Here, we demonstrate the output from simulating the experiment using +microstate. For each participant in the simulation, we assume their ground truth microstate maps to be some group level map plus some noise. Group level maps were arbitrarily chosen and are shown in Figure 3A. Since source flipping is a motivator for using the generalized methodology in +microstate as opposed to classical microstate analysis (Tait and Zhang, 2021), we additionally included source flipping in our individual maps by randomly switching the sign of the maps with 50% probability. An example individual set of ground truth maps are shown in Figure 3B.

**Figure 3:**
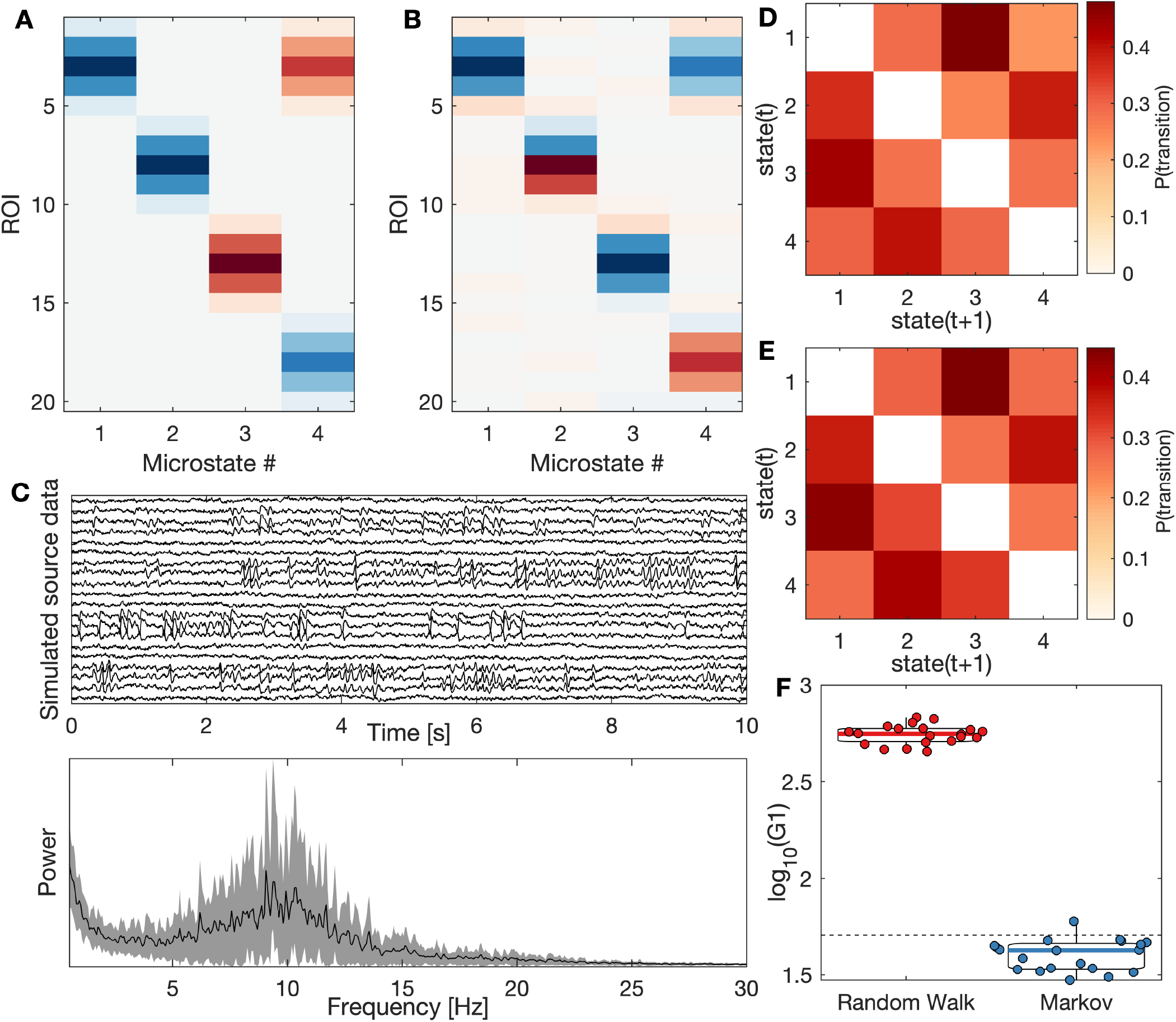
Simulating the experiment described in subsection 3.3. (A) Ground truth group level maps, which are the same for all participants and all conditions. (B) An example individiual ground truth map, which includes random dipole flipping and noise. (C) An example simulation of the source M/EEG data (top) and its power spectrum (bottom). The spectrum shows the mean spectrum across all ROIs, with shaded regions showing standard deviation. (D) Example Markovian syntax matrices for the random walk sequence (i.e. experimental condition 1; top) and Markov sequence (i.e. experimental condition 2; bottom), demonstrating first order Markov properties are consistent across conditions. (E) For the 20 simulated participants, violin plots of the first order Markov statistic (*G*_1_) are shown. *G*_1_ values exceeding the dashed black line are significantly different than a Markovian sequence to *p* < 0.05. While (D) showed first order Markov transitions are the same across experimental conditions, (E) demonstrates that higher order properties of the sequences differ between conditions.

For each participant, a random-walk microstate sequence was simulated and neural dynamics generated as described in (Tait and Zhang, 2021). The resulting simulated source M/EEG and its power spectrum for an example individual is shown in Figure 3. These simulated source data (without ground truth maps or sequences) were stored in a *cohort* structure for later analysis. To generate Markovian surrogate sequences, we used the `stats_markov` function to obtain the Markovian matrix (shown for an example individual without self-transitions in Figure 3D), and subsequently used this matrix to simulate a new Markovian microstate sequence and associated source M/EEG dataset. The simulated M/EEG data for each individual was stored in the same *cohort* object as the data from the random-walk simulations, but using a different condition label to separate the groups. For validation purposes, Figure 3E shows the information theoretical Markov property (von Wegner et al., 2017) for the ground truth random-walk and Markov sequences, showing that (as expected) the random-walk sequences are highly non-Markovian while the Markov sequences are Markovian.

#### 3.3.2 Group-level microstate analysis and comparing conditions

In this section, we will treat the simulated data as though it were experimental source-reconstructed M/EEG and perform group-level microstate analysis. To remove high-frequency noise and slow drifts, we bandpass filtered the simulated data in the range 1-30 Hz (Michel and Koenig, 2018; Tait and Zhang, 2021), and then ran the group level microstate analysis for four states on our *cohort* object.

Figure 4A shows the estimated group-level microstate maps. It is clear that the four estimated maps closely correspond to the group truth group level maps (Figure 3A), excluding polarity. This exclusion of polarity is required for source data to deal with the problem of source flipping (Tait and Zhang, 2021). When aligned to the original group maps using a template matching algorithm, the four estimated maps had map similarities to the original maps between 0.88-0.92, indicating robust estimation of the ground truth microstate maps.

**Figure 4:**
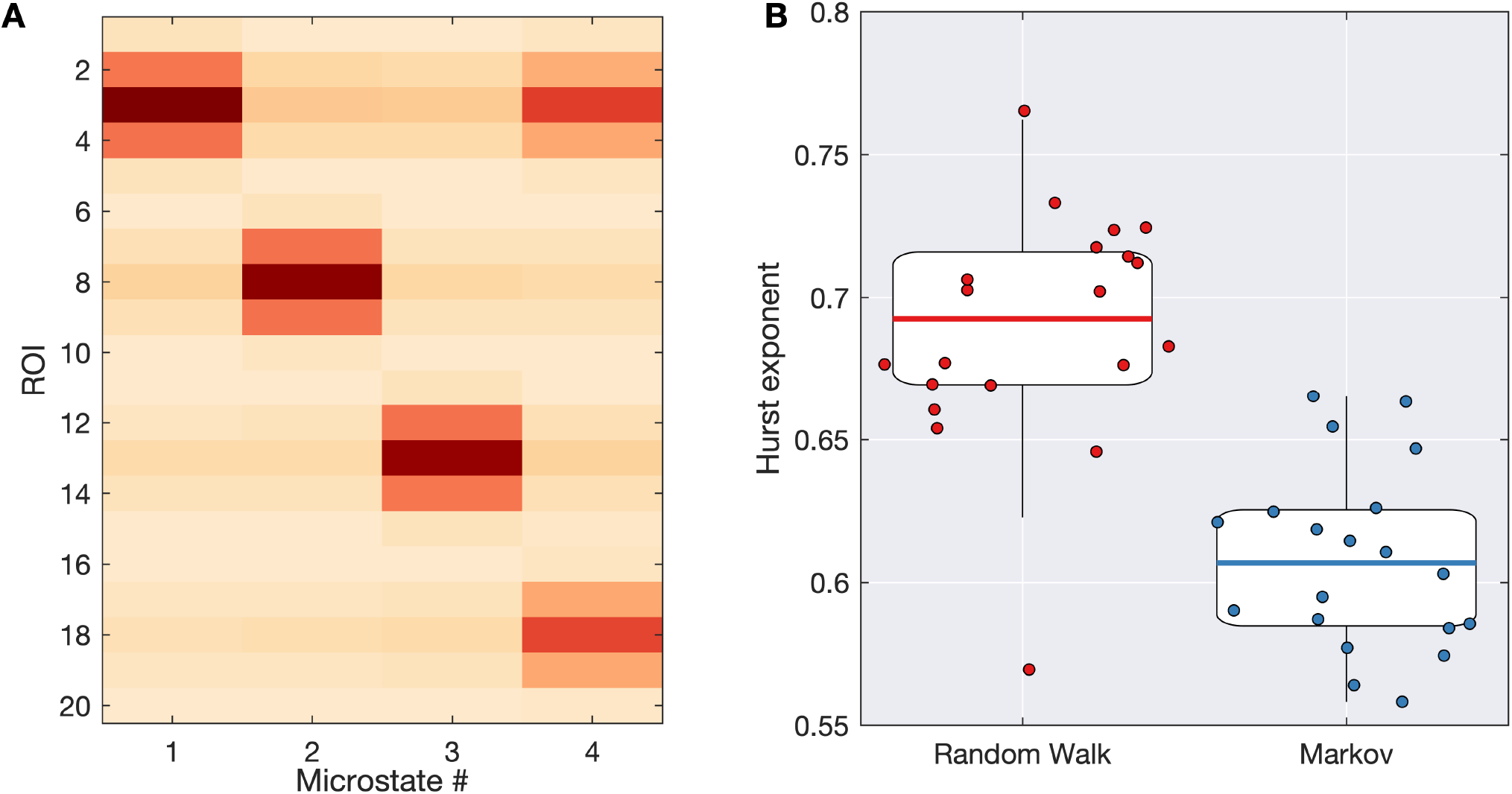
**Group-level microstate analysis** performed on the simulations shown in Figure 3. (A) Estimated group microstate maps. (B) The Hurst exponent of the estimated sequence is significantly lower in condition 2, suggesting fewer long-range correlations than condition 1. Since the underlying simulated sequence for condition 2 was Markovian, this reflects what we should expect.

Next we tested the hypothesis that (estimated) microstate sequences from condition 1 will contain more long-range correlations than sequences from condition 2. To test this hypothesis, we compared the Hurst exponent between conditions, using a Wilcoxon sign-rank test due to the paired experimental design. There was a significantly lower Hurst exponent in the Markov data than the random-walk data (*p* = 2.19 × 10^−4^; Figure 4B), supporting our hypothesis (and validating that the microstate sequences estimated by +microstate reflect statistical properties of the ground truth sequences). Here, we used the Hurst exponent as it is a widely used measure of long-range temporal correlations in microstate sequences, but similar results can be found from studying other measures of non-Markovianity and long-range correlations in +microstate, including the Markov *G*_1_ statistic, the auto-information function, and the microstate sequence complexity.

### 3.4 Source-space MEG microstates response to stimuli

Tait and Zhang (2021) presented a methodology to examine the response of source-space MEG microstates to auditory stimuli, which is implemented in +microstate. In this section, we used +microstate to expand upon this analysis and compare standard vs deviant auditory stimuli. Participants, data acquisition, preprocessing, source-reconstruction, and group-level microstate analysis for the continuous task-based was described in Tait and Zhang (2021). The microstate sequences for all participants and the scripts for the analysis are freely available and included with +microstate.

After importing the continuous task-based data in +microstate, we defined trials from 100 ms prestimulus to 350 ms post-stimulus about each stimulus and concatenated all trials for all participants into a *cohort* object. Tait and Zhang (2021) calculated the *χ*^2^ distance between the histograms of pre-stimulus and post-stimulus microstate probabilities based on the number of samples in each period within each state, demonstrating a significant peak around 100ms following the auditory stimulus. As described in Tait and Zhang (2021), this was a result of an increased likelihood of the microstate containing the auditory cortex. Codes to reproduce the analysis of Tait and Zhang (2021) are included in the tutorials for +microstate.

Here, we expanded upon the analysis of Tait and Zhang (2021) by comparing standard vs deviant stimuli using a similar approach implemented in +microstate. For each sample following the stimulus, we calculated the probability of each microstate (across all participants and trials) for standard stimuli and deviant stimuli. This was used to generate a *χ*^2^ distance between the likelihood of each microstate in the standard vs deviant stimuli at each time point following the stimulus (Figure 5A). By permuting the standard/deviant labels, we performed a cluster-permutation test on *χ*^2^, demonstrating a significant difference in microstate probabilities between 129-250 ms (*p* = 0.0310, cluster permutation test). This time period is consistent with the approximate latency of the auditory mismatch negativity (Rentzsch et al.; Mahadeva Iyer et al., 2017; Nagai et al., 2017). We subsequently plotted the Pearson residuals (Tait and Zhang, 2021) at the sample with peak *χ*^2^ within this time period, representing a histogram of change in microstate probability (shown in Figure 5B). The 2nd microstate, corresponding to the fronto-temporal state ((Tait and Zhang, 2021)) which contains the auditory cortex, was more likely to be active in response to deviant stimuli than standard stimuli. Interestingly, Tait and Zhang (2021) showed that across all stimuli this same microstate is more likely to be active around 100ms post-stimulus than the pre-stimulus period. These results are potentially suggestive of a mechanism by which the fronto-temporal microstate is activated in response to a stimulus (latency approximately 100ms) and remains active for longer during processing of deviant stimuli (latency approximately 130-250 ms).

**Figure 5:**
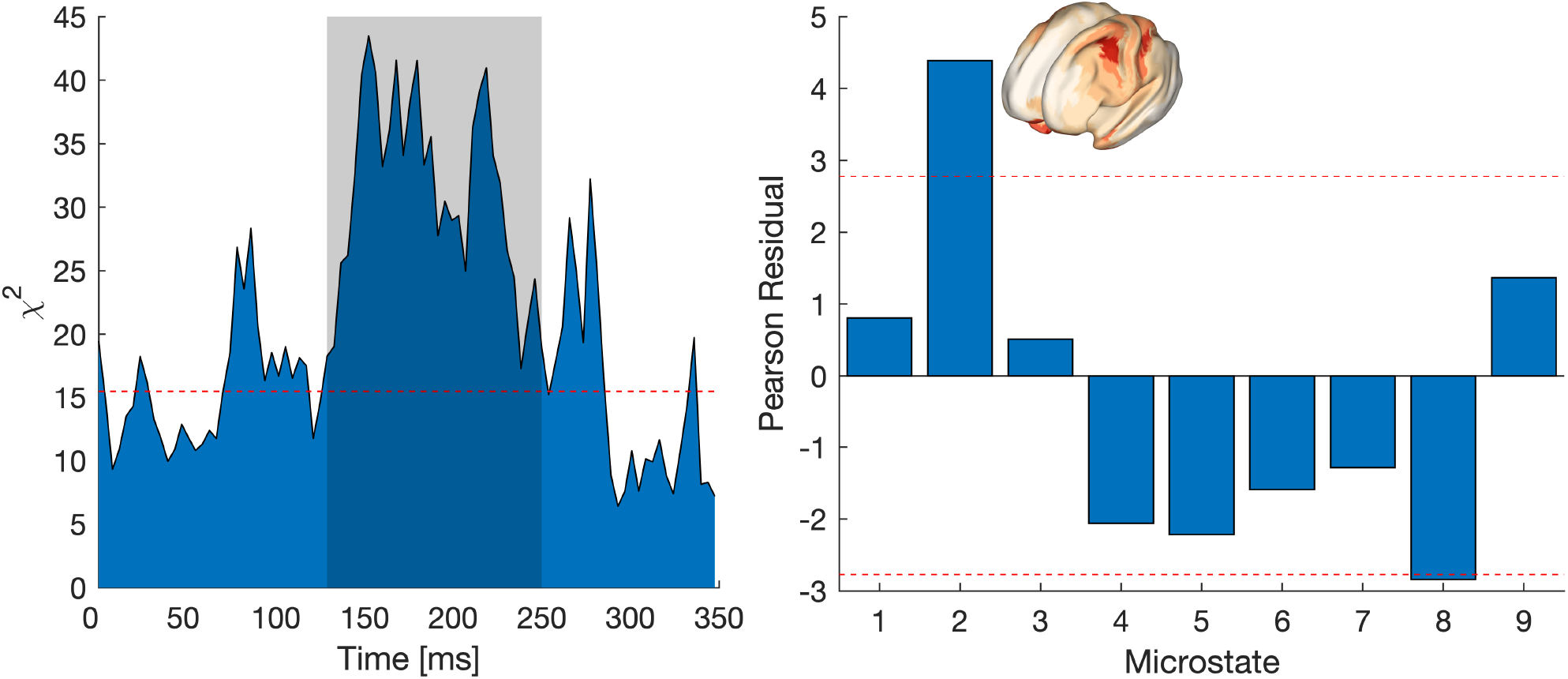
Microstate responses to auditory stimuli. (A) For each sample post-stimulus, the *χ*^2^ distance between the histograms of microstate probabilities for standard and deviant stimuli are shown. The dashed red line corresponds to (uncorrected) *p* < 0.05, which was used as a threshold for cluster permutation analysis. Cluster permutation analysis identified significant differences between 128.9-250.0 ms following the stimulus (shaded region). (B) At peak *χ*^2^ in the significant time period, we plot the Pearson residuals. The dashed red line show Bonferroni corrected *p* < 0.05. Microstate 2 (map for this microstate is inlaid) is significantly more likely for deviant stimuli than standards.

### 3.5 Source space resting-state microstates

Tait and Zhang (2021) used +microstate to examine source-space microstates in resting-state MEG data. Here, we will briefly review their results as an example of the utility of +microstate for anatomical estimation of microstates. Scripts demonstrating how to perform this analysis with +microstate are included in +microstate. These maps explained approximately 60-65% of the variance of the source-reconstructed MEG data, and were robust to new scans.

A key novel feature of the analysis presented by Tait and Zhang (2021) and implemented in +microstate is microstate-segmented functional connectivity. Combining the microstate-segmented functional connectivity patterns with machine learning tools such as multivariate pattern analysis (Treder, 2020), Tait and Zhang (2021) demonstrated that source microstates were associated with distinct patterns of cortical connectivity and can potentially be used as an alternative to arbitrarily chosen sliding windows for studying dynamic functional connectivity states. Study of both microstate activation patterns and microstate synchrony are important for a wholistic understanding of the brain regions involved in generating stable functional brain states, as it is clear that patterns of functional connectivity associated with a microstate are not directly reflective of their activation patterns (Tait and Zhang, 2021).

## 4 Discussion

+microstate is an open-source freely available toolbox for performing microstate analysis in EEG and MEG in sensor and source space. The toolbox includes functionality for pre-processing, microstate clustering analysis, visualization, and statistical analysis at the single-trial and group level. A number of toolboxes for EEG microstate analysis are currently available, including CARTOOL (Brunet et al., 2011) and a plugin for EEGLAB (Poulsen et al., 2018). However, these toolboxes are limited to sensor-space EEG analysis. +microstate is at present the only toolbox for topographic microstate analysis of MEG and source-reconstructed M/EEG data.

It should be highlighted that the focus of +microstate is on the microstate *k*-means algorithm and microstate segmented functional connectivity, and hence there is limited functionality for pre-processing data or statistical analysis. Functions for pre-processing data are limited to re-referencing EEG data, bandpass/bandstop filtering, resampling, orthogonalization and computing amplitude envelopes. More advanced pre-processing steps, notably source-reconstruction, are beyond the scope of the toolbox. Similarly, the only statistical tools implemented in +microstate are non-standard microstate-specific tools such as TANOVA tests. The toolbox was implemented this way by design, since the choice of pipeline for pre-processing and source-reconstruction and the choice of statistical tools used are data and hypothesis dependent. We therefore suggest using +microstate in combination with existing MATLAB toolboxes. For pre-processing and source-reconstruction of M/EEG data, a wide range of toolboxes are available for MATLAB such as Fieldtrip (Oostenveld et al., 2011), Statistical Parametric Mapping (Litvak et al., 2011), EEGLAB (Delorme and Makeig, 2004), and Brainstorm (Tadel et al., 2011). For statistical analysis, in addition to these toolboxes there are options such as the MATLAB Statistics and Machine Learning Toolbox and MVPA-Lite (Treder, 2020). For example, to study source-space MEG microstates,Tait and Zhang (2021) performed data processing and source-reconstruction using Fieldtrip, microstate analysis used +microstate, and statistical analysis used MATLAB functions and MVPA-Lite (Treder, 2020). Other toolboxes such as the Brain Connectivity Toolbox (Rubinov and Sporns, 2010) implemented in MATLAB will likely additionally be useful for analysing properties of the novel microstate-segmented functional networks (Tait and Zhang, 2021) implemented in +microstate. We implemented +microstate in MATLAB on the command line (as opposed to using a graphical user interface), so that the toolbox is easily used in combination with any of these pre-existing MATLAB toolboxes, in contrast to standalone graphical applications for microstate analysis such as CARTOOL.

A number of example use cases were included in this report. The motivations of the use cases were twofold. Firstly, we aimed to demonstrate potential ways that +microstate can be used to analyse (and simulate) data from a range of modalities (sensor-EEG, sensor-MEG, and source-reconstructed data) as well as different cognitive conditions (resting-state and cognitive task). A second motivation for the specific choice of use cases presented here was as validation of the toolbox versus benchmarks and ground truths. In the first example, we analysed resting-state sensor-EEG microstates which have been widely studied in the literature (Khanna et al., 2015; Michel and Koenig, 2018). By reproducing canonical microstate maps and temporal statistics in line with the literature, this example acts as validation that the generalized algorithm used in +microstate gives comparable results to existing EEG microstate literature. In the second example, we performed microstate analysis on sensor-MEG evoked fields between two conditions (congruent and incongruent sentences). Using microstate pipelines implemented in +microstate, we identified topographic differences between conditions at comparable latencies to ERF cluster permutation analysis (Maris and Oostenveld, 2007) implemented in Fieldtrip. We additionally found a microstate map with a topography closely corresponding to the ERF difference map, and found that this map demonstrated significant differences in activation between conditions. This example therefore acts as validation of the sensor-MEG microstate pipeline and topographic ERP/ERF analysis implemented in +microstate. A third example showed simulations of source-space data, and demonstrated that the source-microstate pipeline can reproduce the ground truth microstate maps and group-level differences between conditions. A fourth example contrasted source-reconstructed MEG microstates between standard and deviant auditory stimuli, finding differences at a latency reflecting that of the auditory mismatch negativity. The microstate responsible for these differences include the localization of the auditory mismatch negativity. Finally, we discussed the results of Tait and Zhang (2021) when applied to empirical source-space data (source-reconstructed MEG), particularly highlighting alterations to microstate statistics reflect different cognitive states (resting-state vs auditory task) and the machine learning methods demonstrating associations between microstates and functional networks. Together, these examples provide much evidence for the generalized microstate algorithm implemented in +microstate.

### 4.1 Conclusions

We have presented +microstate, an accessible and freely available toolbox for multi-modal microstate analysis implemented in MATLAB. The aim of this toolbox is to facilitate the use of microstate analysis in a wider range of electrophysiological datasets in future research. Example use cases were given in this toolbox, validating the toolbox and demonstrating use of the toolbox with sensor-EEG, sensor-MEG, and source-reconstructed data across resting- and task-evoked datasets. MATLAB Live Scripts for these examples are included in the toolbox to act as a tutorial to maximise useability and make functional brain microstate analysis an accessible tool for a wider range of researchers.

